# An improved network alignment algorithm of dynamic protein interaction networks of floral development in *Arabidopsis*

**DOI:** 10.1101/2022.11.10.515928

**Authors:** Jiamin Ji, Kailing Zheng, Miao He

## Abstract

To align the dynamic protein-protein interaction networks (DPINs) might help to better understand the molecular network genetic mechanisms of flower development in *Arabidopsis thaliana*. In this paper, the microarray data of flower development in *Arabidopsis* were obtained from the gene expression omnibus (GEO) database. The active expressed genes at each time point were screened using σ method. The DPIN at each time point were reconstructed using search tool for the retrieval of interacting genes/proteins (STRING) and then visualized with Cytoscape, and enrichment analysis implemented using clusterProfiler package of the R language. We improved NetCoffee 2, a multi-time, multidimensional, and multi-level network alignment algorithm, NetCoffee 2+, was developed for floral development of A. thaliana. In this paper, a network alignment algorithm applied to *A. thaliana*, NetCoffee 2+, was developed, and the key proteins and protein complexes during floral development in *A. thaliana* were mined, with new potential ABCDE-like genes discovered.

## 1 Introduction

### 1.1 Problem of flower development

Flowering is the process in transformation from the vegetative growth to the reproductive growth in angiosperms. Flower development in plant is the popular scientific issue in plant physiology, molecular biology, genetics and developmental biology.

The most important achievements of molecular genetics on flower development are mainly due to *Arabidopsis thaliana*. In the early 1990s, genetic studies using floral organ mutants in *Arabidopsis* and *Antirrhinum majus*, representing mutations in mainly MADS box transcription factor genes, led to the establishment of the robust ‘ABC model’ for floral organ formation [1]. Subsequently, the D-class genes and E-class genes, further refining types of genes involved in flower development, had been reported [2-3]. The five types of genes except the A-class gene *APETALA2* (*AP2*) belong to the MADS-box gene; among them, the E-class genes can bridge the MADS-box genes to form function complexes [4]. Combined with the further improved pattern of flower organ development, the widely accepted ABCDE model had been proposed (see **Figure 1**). The floral quartet model elucidated the regulation pattern of flower organ development in protein level [5]. It was assumed that different combinations of four homeomorphic gene products or protein complexes (PCs) determine the morphological characteristics of different floral organs. These PCs bind to the target gene promoter to activate or inhibit the function of genes related to different floral organ characteristics.

**Figure 1.**
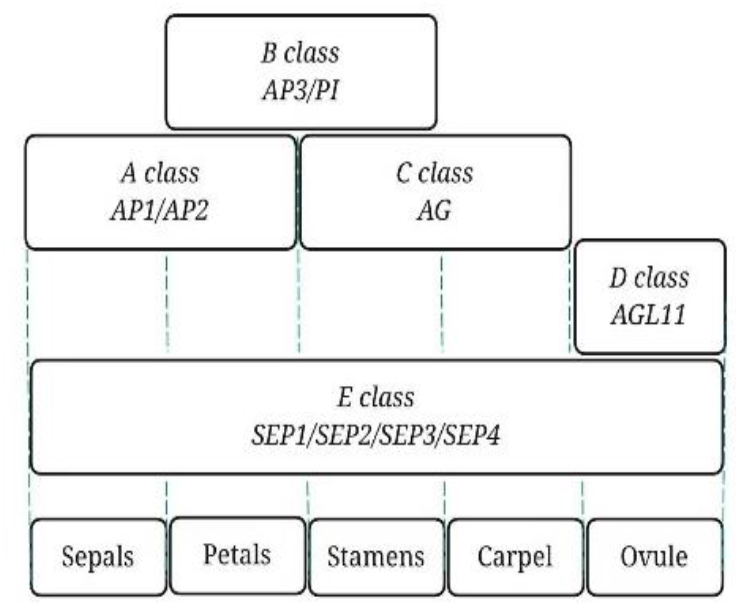
ABCDE mode of flower development

### 1.2 Biomolecule network alignment

Interactions between proteins perform important biological functions. Therefore, deciphering the connectivity pattern or topology of protein-protein interaction networks (PPINs) is critical for clarifying the basic functions of cell. By performing global alignment between PPINs, annotations can be transferred between PPINs, thereby screening out the basic constituent elements and evolutionarily conserved parts of PPINs. Combining interacting protein topology and sequence similarity, Kelley et al. had aligned two PPINs, demonstrating that these PPINs of *Saccharomyces cerevisiae* and *Helicobacter pylori* harbor a large number of evolutionarily conserved pathways, and that many pathways appear to be replicated and specialized in yeast [6]. An algorithm based solely on network topology, Kuchaiev et al. aligned the PPINs of *Saccharomyces cerevisiae* and *Homo sapiens*, showing that even very distinct species share a surprising number of similar regions [7]. Bandyopadhyay et al. aligned the PPINs of two species by assigning proteins to clusters of sequence homology [8]. Network analysis could also be used for refining sequence-based homology searches provides key sources of information. By integrating different cost functions and alignment strategies, Faisal et al. combined a novel PPIN alignment method and used this method to convert aging-related information from correctly annotated model species S*accharomyces cerevisiae, Drosophila melanogaster* and helminth to poorly annotated *Homo sapiens*, resulting in new insights into human ageing-related knowledge that could complement existing knowledge about ageing that is currently primarily obtained through sequence alignment [9].

We hope to align the dynamic protein-protein interaction networks (DPINs) which might help to better understand the molecular network genetic mode of flower development in *Arabidopsis thaliana*.

## 2 Data and method

### 2.1 Data

The microarray data were obtained from the Gene Expression Omnibus (GEO) database, GSE64581 contains a total of 42 microarrays at 14 time points during flower development in *Arabidopsis*. The data were treated with dexamethasone (10 *μM*) at different time points (0d, 0.5d, 1d, 1.5d, 2d, 2.5d, 3d, 3.5d, 4d, 4.5d, 5d, 7d, 9d, 11d and 13d) which were taken from three samples.

### 2.2 Method

#### 2.2.1 Definition

For a given K networks (or subgraphs), *G*_*n*_ = (*V*_*n*_,*E*_*n*_), 1 ≤ *n* ≤ *k*, where *V*_*n*_ and *E*_*n*_ represent the set of nodes (proteins) and edges (PPIs) of the network *G*_*n*_, respectively. The goal of network alignment is to find one-to-one (or many-to-many) correspondences *M*, i.e. a set of node pairs (*u, v*)∈*Vi* ×*Vj* ((*Su, Sv*) ⊂ *Vi* ×*Vj*),*i* ≠ *j*,1 ≤ *i,j* ≤ *K*, where *u* and *v* (i.e. *S*_*u*_ and *S*_*v*_) are well conserved in sequence and topology.

#### 2.2.2 Uniquely expressed proteins at each time point

At different time points of *Arabidopsis* flower development, different proteins and protein complexes perform different functions. *Arabidopsis* flower development is a continuous process. From a molecular point of view, most genes are present continuously during *Arabidopsis* flower development, but each time point has its own uniquely expressed proteins. The uniquely expressed proteins at each time point may perform important biological functions, so this paper screened the functions of the uniquely expressed proteins at each time point. By looking for uniquely expressed proteins at each time point, combined with literature, analyze the role of *Arabidopsis* flower development at the level of individual proteins.

#### 2.2.3 DPIN alignment

DPIN alignment can help identify evolutionarily conserved pathways or protein complexes that may be structurally or functionally important. It is possible to speculate on the role of proteins of unknown function, or speculate on proteins that perform the same function in different spaces and times.

In this paper, we developed a multi-time, multi-level and multi-dimensional network alignment algorithm NetCoffee2+.

The main principle of NetCoffee2+ is as follows. For a given network *G* = (*V,E*), *V* = (*v*_1_,*v*_2_,…,*v*_*n*_), *v*_*i*_ ∈*V*, define a six fold eigenvector (*α,β,γ,δ,θ,ε*), representing the locally connected properties of each node in *V*. Without loss of generality, the adjacency matrix that defines the network *G* is *M*_*n*×*n*_, because *M* is a real symmetric matrix, there must be a normalized eigenvector *K* = (*k*_1_,*k*_2_,…,*k*_*n*_). In other words, *K* is the eigenvector of the largest eigenvalue. Define the centrality of the eigenvector of a node *V*_*i*_ as *k*_*i*_, that is, the centrality of the node i is equal to the element i in the eigenvector, the larger *k*_*i*_ is, the more important the node *V*_*i*_ is, denoted *k*_*i*_ as *α*; define the set of adjacent nodes to the node *V*_*i*_ as *N*_*v*_, choose | *N*_*v*_ | as *β*. The sum of the centrality of all nodes in the *N*_*v*_ definition 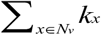 is *γ*; the set of nodes that are two steps away from the node *V*_*i*_ and have no direct relationship with the node *V*_*i*_ is defined as 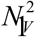, the selection 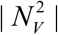 is *δ*; use the formula 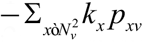 to calculate the fifth element, which *p*_*xv*_ represents the number of the shortest distance from *x* to *v*, and record for*θ*. To align the DPINs at adjacent time points, the unknown protein functions are predicted, and the PPI roles are analyzed.

#### 2.2.4 Protein complex mining

In order to mine vital proteins and PCs in DPIN, we use dendritic cell algorithm (DCA) as the supplement of NetCoffee2+, which is applied to DPINs alignment. DCA can be used to predict unknown protein functions, and analyze at PCs level.

## 3 Results

Combined with literature screening of vital proteins in the whole process DPINs of flower development, a total of 217 key proteins were obtained from uniquely expressed proteins at each time point and aligned proteins at adjacent time points.

The genes encoding these proteins were clustered with the A-, B-, C-, D-, E-class genes. The results showed that, *HTR11, UPB1, HWI1, SNS1, QUL1, MCC1, PUB12, CML5, PRK4, GDPD6, LSM1A, TFIIB2* are the potential floral function genes, in which *TRO, 4CL8, CRK29, RHS16, LOX4, OPS, CSI1, CDC48B, CVY1, EOL1, CSTF64, IWS1*, *CFM4, FUF1* and *DGK5* are the potential A-class genes, and *GATA3, EDE1, MHF2, PRIN2, FEI1* and *PCR2* are the potential C-class genes, *NLP1, ARI10, RSL1, NSP4, NSP3* and *HBI1* are the potential A-class or D-class genes (see **Figure 2**).

According to the DPIN scale at each time point, it can be found that in the early stage of flower development, DPIN contains more active proteins, and the networks maintain stable scales, genes related to flower development are activated and gradually join the networks. This period is from day 0 to 2.5, covering stages 1 to 3 of flower development. During this period, the flower primordia begins to form and develop, and plants mainly carry out photosynthesis and carbohydrate metabolism to provide energy for subsequent development processes. Among the key proteins in this period, CVY1, AMP1, and PKT4 are involved in regulating the transition from vegetative growth to reproductive growth in *Arabidopsis*; AtERF014 is responsive to pathogen infection and defense signaling g hormones; CMT1, LPA19 are involved in chloroplast development, photosynthetic activity, and PSII complexes PILS6 may be related to temperature sensing; HSP70, Cpn60β, MCTP1, etc. are involved in regulating the flowering time of *Arabidopsis thaliana*, which may affect the function of flowering integrators in different ways during the flowering transition. FUF1 and KAT1 are involved in the transduction of phytohormone signals and regulate the growth of plants. PC0-1 regulates plant response to light, participates in photosynthesis in *Arabidopsis* flower buds, and plays a role in the initiation of flower development. PC0-3 are involved in plant responses to environmental changes and are related to bioremediation. There is synergy between PC0-4 and PC0-1 and PC0-3. PC1.5-2 is involved in the formation of photosystem, and PC1.5-4 is involved in wound repair. PC2.0-5 is involved in the detoxification process of cellular methylglyoxal (MG). In the middle stage of flower development, the number of actively expressed proteins decreased. It is speculated that during this time period, the proteins involved in the decision process of flower development have been expressed, and the genes involved in the regulation of flower development process began to be expressed. This period is the 3rd to 7th day of flower development, covering the 3rd to 7th stage of flower development. During this period, the calyx primordium, petal primordium, stamen primordium, and pistil primordium gradually formed and continued to develop in turn. Plants mainly carry out physiological processes such as sugar metabolism and fatty acid metabolism. Plants are in a state of rapid differentiation, and various structures of flower organs are gradually formed. Among the key genes in this period, there is a correlation between BCATs genes and plant flowering time. The dynamic response of starch polymers to day length is regulated by CO through modifying expression of GBSS1 during the flowering transition period. The flowering-inducing function may not be limited to regulating FT expression, it may also play a vital role in controlling the metabolic components that provide resources for floral transformation. SAUR is involved in auxin signal transduction; ROXY1/2, MHF2, DGK5, MCC1, ABCG9/ ABCG31 are involved in regulating anther development and regulating anther’s adaptation to the environment; in addition, ROXY1/2 is also involved in controlling the development of sepals and petals. PC3.5-2 is involved in the maintenance of copper homeostasis and regulates flowering time. In the later stage of flower development, the number of actively expressed proteins increased sharply. It is speculated that during this period, flower organs have basically formed, mainly involving complex physiological processes such as pollen maturation, stigma development, and insemination, and more proteins need to be mobilized for regulation. This period is the 9th to 13th day, covering the 9th to 12th stage of flower development. During this period, the stigma appears above the pistil, and the petals continue to grow until they are level with the stamens. Plants mainly undergo physiological processes such as mitosis, meiosis, and DNA replication. Plants are in a stage of rapid growth, and cells proliferate rapidly. Among the key proteins in this period, INA2 is related to heat tolerance in the process of plant reproduction and development, ISTL1 is essential in the process of pollen development; NOXY7 is involved in plant immunity; BNQ2 is involved in the regulation of flowering; RLK, MCU2, BZIP34 are involved in pollen tube germination and growth; DRMY1 is involved in abscisic acid signaling and plays an important role in organ development; UXTI/2/3 co-regulate xylan biosynthesis, and pollination is impaired in triple mutants, presumably caused by changes in cell wall composition. Reduced filament growth and anther dehiscence, the filament growth length is lower than the stigma, resulting in pollen grains not falling on the stigma normally for pollination. KCS4 is involved in the biosynthesis of plant ultralong fatty acids; in contrast to major advances in photosynthesis or storage organs, the floral lipidome is an unexplored frontier in plant lipid research. However, substantial evidence from recent molecular biology studies suggests that lipids play a crucial role in coordinating flower development, rather than as inert end-products of metabolism. PC9.0-2 is functionally similar to PC9.0-3, involved in the occurrence and development of female gametes, and is redundant during flower development. PC11.0-1 is involved in embryonic development. PC11.0-2 is involved in DNA repair. PC11.0-3 is involved in stamen development in *Arabidopsis*. PC11.0-1/2/3/4 may combine with DNA, RNA and other substances, cooperate with each other, and participate in regulation flower development of organ phenotype. PC13.0-1 is involved in nucleobase, nucleoside, nucleotide and nucleic acid metabolism (see **Figure 2**).

**Figure 2.**
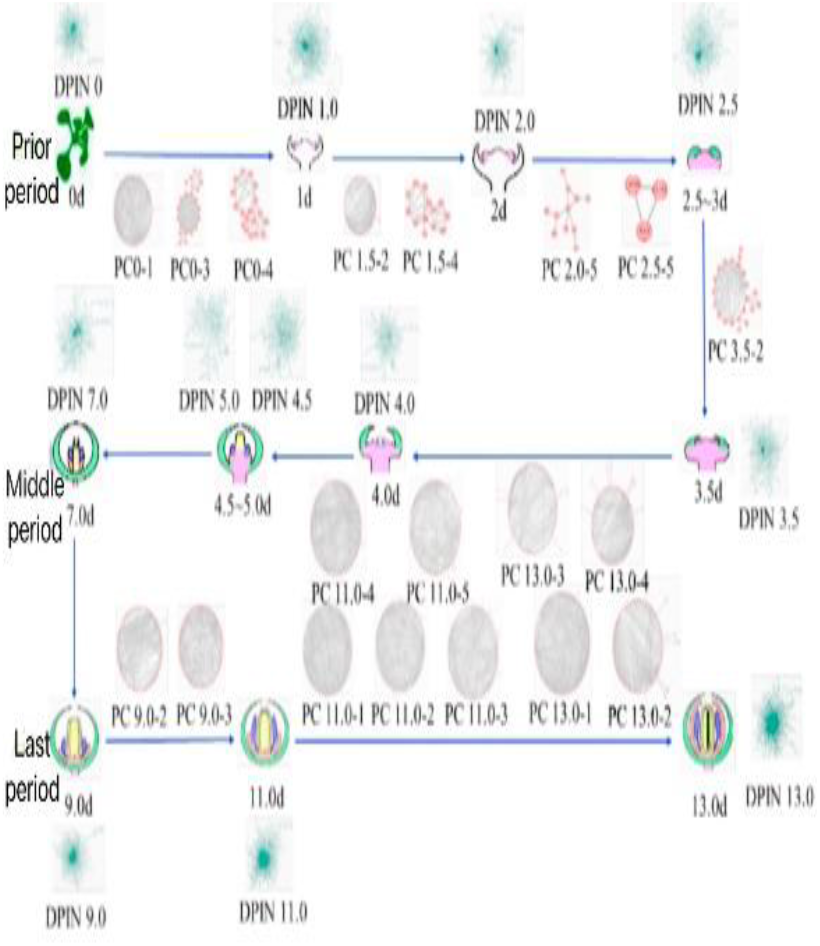
Key protein complexes throughout the whole process of flower development.

This research was funded by the General Program of the National Natural Science Foundation of China (Approval Number: 31870348).

